# Targeted mRNA demethylation using an engineered dCas13b-ALKBH5 fusion protein

**DOI:** 10.1101/614859

**Authors:** Jiexin Li, Zhuojia Chen, Feng Chen, Yuyi Ling, Yanxi Peng, Nan Luo, Hongsheng Wang

## Abstract

Studies on biological functions of *N*^6^-methyladenosine (m^6^A) modification in mRNA have sprung up in recent years. Here we construct and characterize a CRISPR-Cas13b-based tool for the first time that targeted m^6^A methylation of mRNA by fusing the catalytically dead Type VI-B Cas13 enzyme from Prevotella sp.P5-125 (dPspCas13b) with the m^6^A demethylase ALKBH5, which is named as dm^6^ACRISPR. Subsequently, such system is shown to specific demethylase the m^6^A of target mRNA such as CYB5A to increase its mRNA stability. In addition, the dm^6^ACRISPR system appeared to afford efficient demethylation of the target genes with tenuous off-target effects. Together, we provide a programmable and *in vivo* manipulation tool to study mRNA modification and its potential biological functions of specific gene.

## INTRODUCTION

As the prominent dynamic mRNA modification, *N*^6^-methyladenosine (m^6^A) has been identified since the 1970s ^1^, while thousands of RNA transcripts contain m^6^A modifications with unique distribution patterns as well ^2^. m^6^A was governed by methyltransferase complex (“writers”), demethylases (“erasers”) and RNA-binding proteins (‘readers’) ^3^. It can regulate the splicing, translation, and decay rates of mRNA to affect protein production. m^6^A modifications have been reported to regulate various biological processes such as heat shock response ^4^, ultraviolet-induced DNA damage response ^5^, maternal mRNA clearance ^6^, neuronal functions and sex determination ^7^, cortical neurogenesis ^8^, haematopoietic stem and progenitor cell specification ^9^, and T cells homeostasis ^10^. However, whether m^6^A modification could regulate the mRNA processing of metabolic-related genes and affect metabolism remains unknown.

Current methods of manipulating RNA methylation are primarily based on modulating the expression of RNA methyltransferases (Mettl3/Mettl14/WTAP complex) or demethylase (FTO and/or ALKBH5) via bioengineering methods, which cause broad epigenetic changes and activation of endogenous retroviruses ^9, 11-14^. Recently, flavin mononucleotide was identified as a cell active artificial m^6^A RNA demethylase, which provided a potential and powerful small molecule for RNA demethylation ^15^. However, as these methods and reagents demethylate transcriptome globally, it is difficult to study the effect of specific RNA methylation events, and there is the risk of side effects in therapeutic use. A lack of technologies for targeted manipulation of RNA methylation has also hindered study of the correlation between locus-specific RNA methylation and gene expression. Generating an easy-approached RNA demethylation has great potential utility for dissecting the role of RNA methylation in multiple biological processes.

The recently discovered CRISPR/Cas system has provided a great potential way to study the endogenous functions and dynamic variations of nucleic acids^16-20^. The nuclease-inactive DNA-targeting Cas9 (dCas9) has been used to deliver the interested proteins to the target sites via fusion protein form ^16-20^. The discovery of RNA-targeting Cas proteins Cas13 opens the door for targeting the dynamic of endogenous RNA transcripts ^21^. Similar to Cas9, mutation of nuclease domains of Cas13 can also generate a catalytically dead enzyme that retains RNA binding affinity (dCas13). The nuclease-inactivate “dead” version of LwaCas13a (dLwaCas13a) can visualize the endogenous RNA via fusion with EGFP^22^. The fusion with A-to-I RNA editing enzymes ADAR1 and ADAR2 of PspCas13b can direct edit RNA at endogenous RNA sites ^23^. All these studies indicated that Cas13-based systems can act as a powerful tool to investigate the endogenous RNA regulation.

The fused enzymes and dCas9 can modify the epigenetic properties such as DNA methylation and histone methylation/acetylation status ^17, 19, 24-28^. Further, recent study showed that dCas13b-m^6^A reader tools can target specific transcripts of interest to regulate the mRNA translation and degradation ^29^. In the present study, we constructed and characterized a CRISPR-Cas13b-based tool for the first time that targeted m^6^A methylation of mRNA by fusing the catalytically dead Type VI-B Cas13 enzyme from Prevotella sp.P5–125 (dPspCas13b) with the m^6^A demethylase ALKBH5 via a six amino-acid (GSGGGG) linker. This tool combines the ease of Cas13 protein using short complementary single guide RNA (sgRNA) with the already established ALKBH5 activity in fusion constructs ^30^. We demonstrate that our constructs maintain the previously reported effects on mRNA demethylation and increase the stability of targets, which is named as **dm**^**6**^**ACRISPR**. We go on to show that the fusion proteins can decrease the expression of endogenous transcripts in an ALKBH5-dependent manner, suggesting that this tool may find utility for biotechnological applications. Together, this work provides a programmable and *in vivo* manipulation tool to study mRNA modification and its potential biological functions of specific genes.

## RESULTS

### 1. Design of dm^6^ACRSIPR for targeted RNA demethylation system

To provide a simple tool to study RNA modifications, the dCas13b-ALKBH5 fusion protein was constructed by addition of ALKBH5 to the C–terminus of the inactive Cas13b (dCas13b) with a six amino-acid (GSGGGG) linker or by prior of ALKBH5 to the N-terminus of dCas13b with the same linker (Figure 1 A and B). Meanwhile, U6 promoter-driven gRNA transcription system was cloned into the pC0043-PspCas13b gRNA backbone by use of BbsI ^23^. Both dCas13b-ALKBH5 and ALKBH5-dCas13b fusions were efficiently expressed in cells as illustrated by Western blotting (Figure S1A). In HEK293T cells, the transfection of both fusion proteins can significantly decrease global m^6^A of mRNA, suggesting the *in vivo* demethylation function of ALKBH5 fusion proteins (Figure S1 B). However, the efficiency of both dCas13b-ALKBH5 and ALKBH5-dCas13b was less than that of ALKBH5 (Figure S1 B), which might be due to the increase of steric hindrance of fusion proteins that impairs their catalytic activities. Presence of NES (nuclear export signal) could export fusion proteins in the cytoplasm as compared to that of ALKBH5 which mainly located at the nuclei, which was confirmed by immunocytochemistry using anti-ALKBH5 antibody (Figure S1 C).

**Figure 1.**
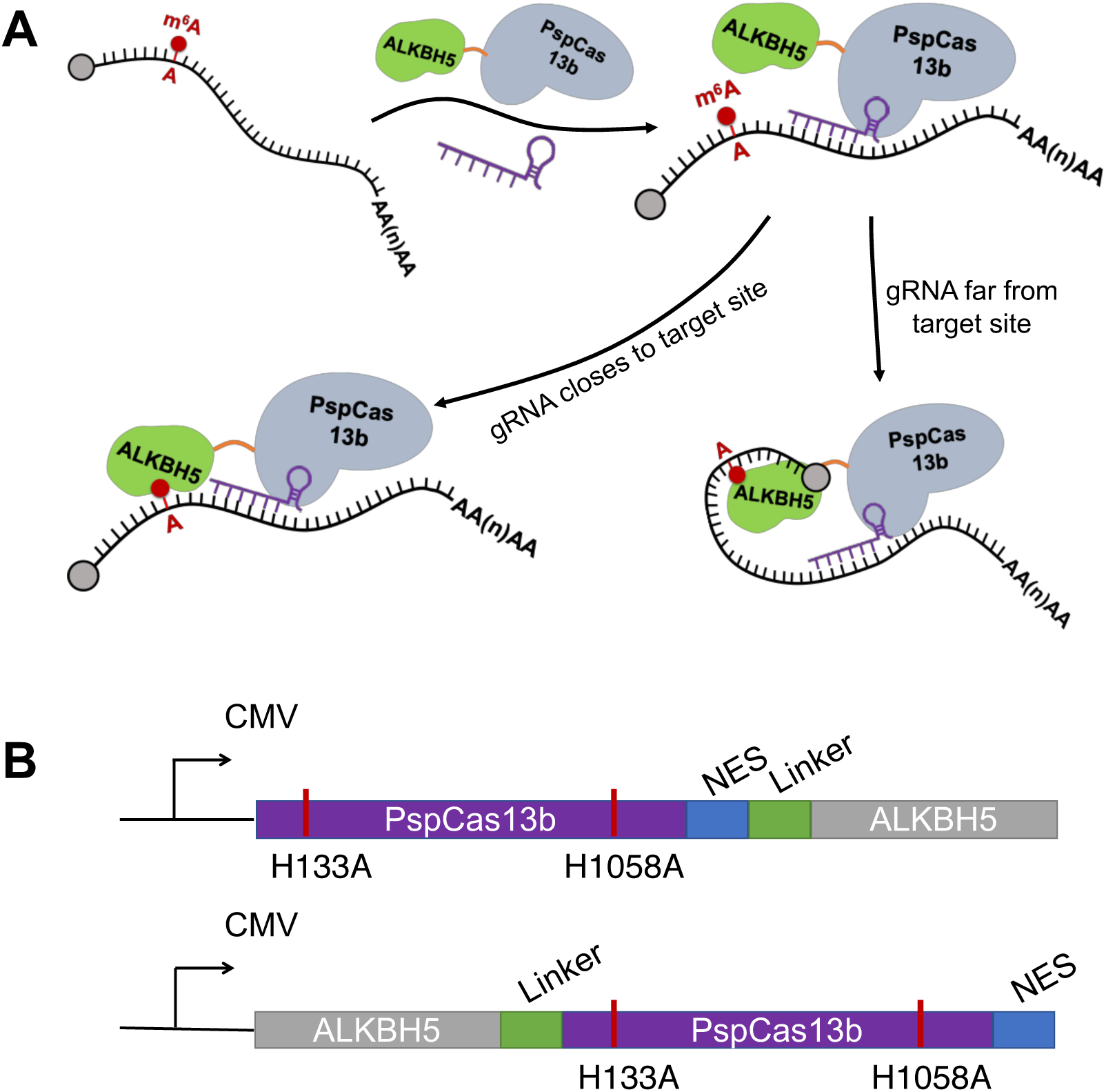
Design of CRISPR-dCas13b-based targeted RNA demethylation system. (A) General overview of site-specific RNA targeting using dCas13b-guided fusion proteins; (B) Schematic illustration of the domain organization of the dCas13b-ALKBH5 expression cassette.

### 2. The dm^6^ACRISPR induced m^6^A demethylation at CDS increases mRNA stability

To test the utility of dCas13b-ALKBH5 for targeted m^6^A methylation, we selected potential candidate transcripts according to the following rules: 1) The RFKM of transcript is greater than 300 in m^6^A-seq according to our previous study; and 2) Only one m^6^A peak site was identified for one transcript. For those transcripts bearing m^6^A variated in cells undergoing EMT (in our previous study) ^31^, they are preferentially selected due to the more dynamic variation of m^6^A. Finally, the transcript of CYB5A was chosen to investigate the effects of dCas13b mediated demethylation system (Figure 2 A and Figure S2A). m^6^A-RIP-seq data showed that unique m^6^A peak exists in CYB5A CDS region at 48 A with the motif of GGA (Figure S2A). The m^6^A modification in mRNA of CYB5A was ensured by m^6^A-RIP-qPCR in both HeLa and HEK293T cells (Figure 2B). Further, the m^6^A site in CDS region of CYB5A at 48 A was verified by “SELECT” method ^32^ in HeLa and HEK293T cells as well as their corresponding Mettl3 knocked down cells (Figure 2C), which also confirmed that the m^6^A at 48A of CYB5A CDS is exist and reversible.

**Figure 2.**
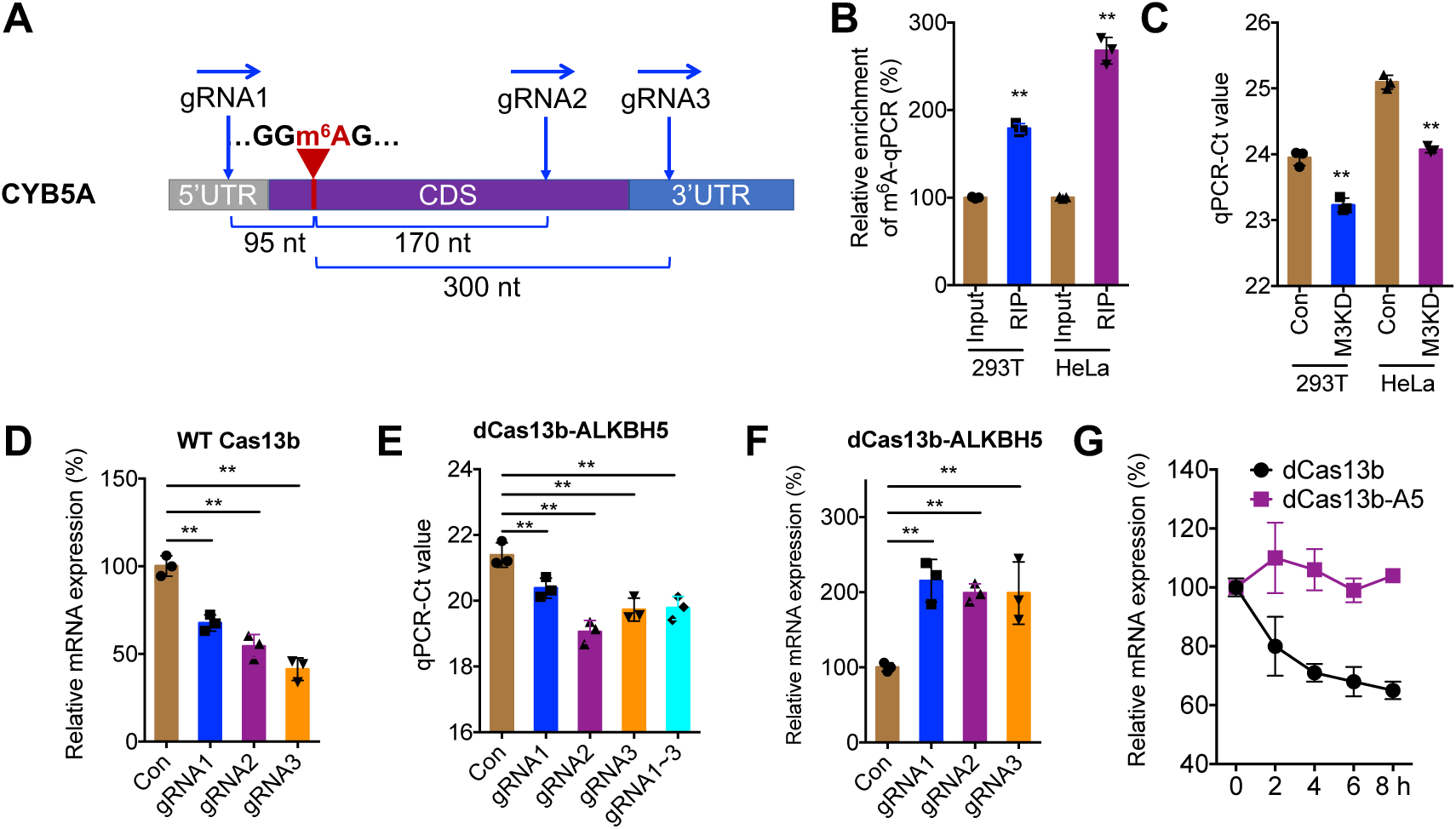
The dm^6^ACRISPR induced m^6^A demethylation at CDS increases mRNA stability. (A) Schematic representation of positions of m^6^A site within CYB5A mRNA and the regions targeted by three gRNAs, respectively; (B) m^6^A RIP-qPCR analysis of CYB5A mRNA in HeLa and HEK293T cells; (C) The threshold cycle (CT) of qPCR showing SELECT results for detecting m^6^A site in CDS region of CYB5A at 48 A in HeLa and HEK293T cells as compared to their corresponding Mettl3 knocked down cells; (D) The expression of CYB5A in HEK293T cells transfected with wild type Cas13b and gRNAs for 24 h; (E) The threshold cycle (CT) of qPCR showing SELECT results for detecting m^6^A site in CDS region of CYB5A at 48 A in HEK293T cells transfected with dCas13b-ALKBH5 combined with gRNA control or gRNA1/2/3, respectively, for 24 h; (F) The mRNA expression of CYB5A in HEK293T cells transfected with dCas13b-ALKBH5 combined with gRNA control or gRNA1/2/3, respectively, for 24 h; (G) HER293T cells were pre-transfected with dCas13b or dCas13b-ALKBH5 with gRNA2 for CYB5A for 24 h and then further further treated with Act-D for the indicated times, and mRNA expression of CYB5A was measured by qRT-PCR. Data are presented as the mean ± SD from three independent experiments. ** p<0.01 compared with control.

The mRNA of CYB5A was targeted by three gRNAs at distinct positions, around the m^6^A site (Figure 2 A). To test the efficiency of gRNAs, we checked the mRNA expression of CYB5A in cells transfected with gRNAs and wild type Cas13b, which cleaves targeted mRNA. Our data showed that all three gRNAs combined with wild type Cas13b significantly decreased the mRNA levels of CYB5A (Figure 2 D), suggesting that all three gRNAs combined with wild type of Cas13b can work efficiently *in vivo*. While transfection of gRNAs alone (Figure S2 B) or gRNAs combined with dCas13b (Figure S2 C) had no effect on mRNA expression of CYB5A.

To minimize the off-target effects of overexpressing artifacts^33^, we optimized the transfection amount of fusion proteins and compared their effects on mRNA expression of interests compared with that of ALKBH5 and control groups. The data showed that the capability of dCas13b-ALKBH5 induced upregulation of targets was at least 6-fold less than that of ALKBH5 (Figure S2 D). Therefore, 1.5 μg dCas13b-ALKBH5 plasmid per 1 × 10^6^ cells was used for further studies due to the non-significant effect on mRNA expression of targets.

We then checked the effect of gRNAs and dCas13b-ALKBH5 on m^6^A modification of CYB5A. SELECT showed that all three gRNAs combined with dCas13b-ALKBH5 can significantly decrease the m^6^A levels of targeted site (Figure 2 E). The strongest demethylation on CYB5A CDS was observed with gRNA2, which targets a ∼170 nt downstream region from the m^6^A site and resulted in 80.2 ± 3.6% demethylation (2-ΔCt method). The dCas13b-ALKBH5 mediated demethylation of CYB5A was further confirmed by m^6^A-RIP-qPCR (Figure S2 E). Intriguingly, gRNAs and dCas13b-ALKBH5 transfection leaded a significant upregulation of CYB5A mRNA (Figure 2 F). To investigate whether dCas13b-ALKBH5 induced upregulation of CYB5A was due to m^6^A mediated mRNA decay, we compared the effects of dCAS13b-ALKBH5 or dCas13b with gRNA2 on CYB5A mRNA half-life. Results showed that targeted demethylation of CYB5A can significantly stabilize its mRNA (Figure 2 G), suggesting that dm^6^ACRISPR can increase the mRNA stability via demethylating m^6^A at CDS in the case of CYB5A.

### 3. Specificity and non-additive effects of dm^6^ACRISPR on m^6^A demethylation

Having shown the efficient demethylation of the target sites, the question appeared if off-target RNA demethylation is introduced at additional transcriptome locations. To check this, we predicted off-target gRNA binding sites for the gRNAs of CYB5A by BLASTN using “somewhat similar sequences” ^**27**, **34**^. PRSS56, GALR1, and INPP4A had the highest matches for gRNA1/2/3 of CYB5A, with the matches of 86%, 92%, and 77%, respectively. The methylation statuses of these off-target loci were measured after transfected with dCas13b-ALKBH5 and their corresponding gRNAs. m^6^A-RIP-PCR results showed that there was no significant effect on the methylation levels of PRSS56, GALR1, or INPP4A (Figure 3 A∼C), although two of them showed a slight demethylation increase (4.8% and 6.4 % decrease of methylation on GALR1 and INPP4A, respectively). Comparing to the 80.2% and 69.4 % demethylation efficiency on CYB5A mRNA using gRNA2^CYB5A^ and gRNA3^CTB5A^, the off-target effect on tested transcripts was limited.

**Figure 3.**
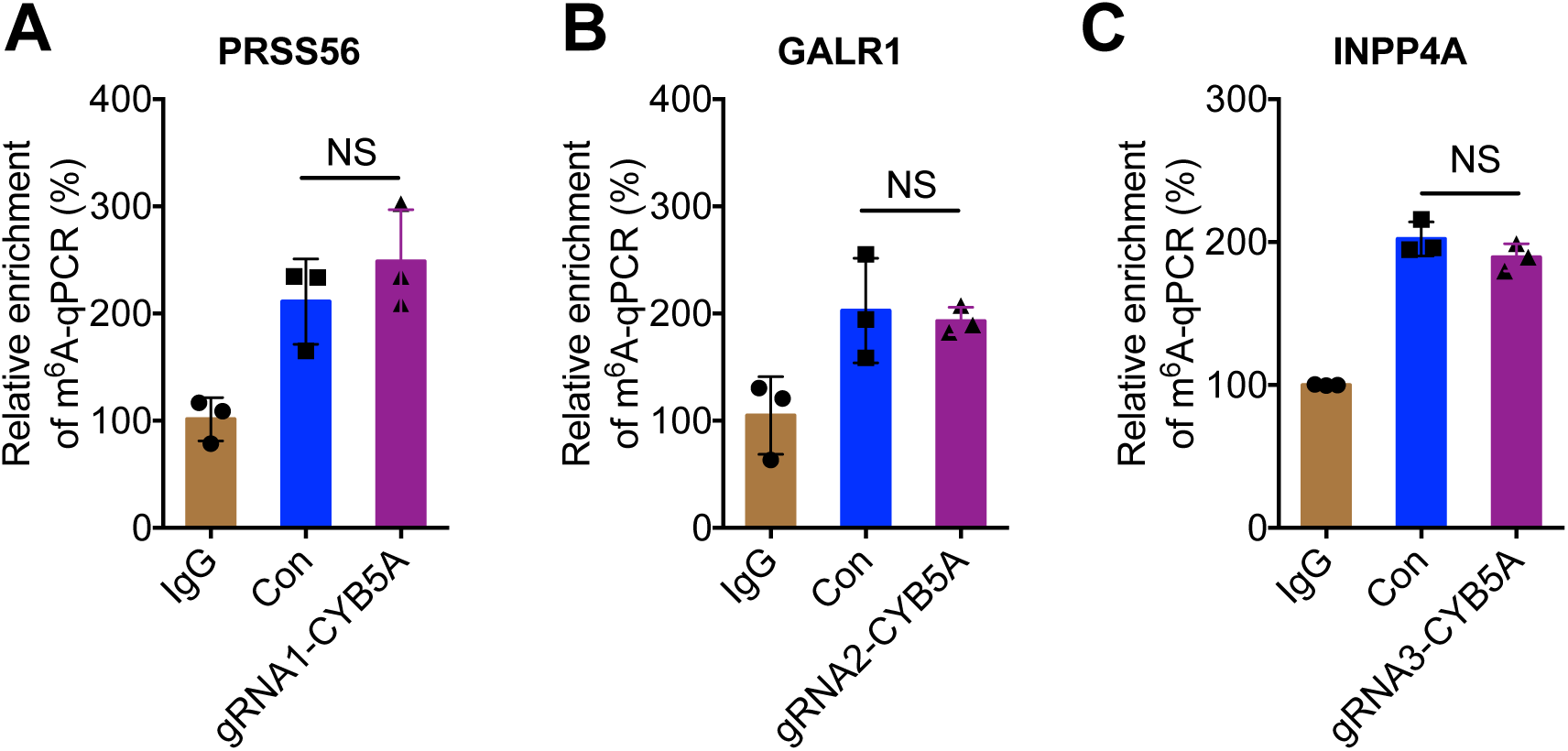
The specificity of dm^6^ACRISPR. Cells were transfected with dCas13b-ALKBH5 and gRNA1/2/3 of CYB5A, respectively, for 24 h, then the m^6^A of PRSS56 (A), GALR1 (B), or INPP4A (C) was measured by m^6^A -RIP-qPCR analysis.

Subsequently, we tested whether co-targeting same loci with multiple gRNAs could further enhance the demethylation levels. However, targeting all three gRNAs to the CYB5A CDS did not increase the demethylation efficiency (68.3 ± 3.5%, Figure 2 E) when compared to single gRNA targeting experiments (e.g. gRNA2, 80.2 ± 3.6%). It suggested that there is non-additive effect of the dm^6^ACRISPR system for RNA demethylation.

## DISCUSSION

In summary, we conclude that single gRNAs co-transfected with dCas13b–ALKBH5, named as dm^6^ACRISPR, can cause robust mRNA demethylation for the specific transcript with limited off-target effects. Specifically, the dm^6^ACRISPR mediated demethylation of CDS can increase the mRNA stability of CYB5A. The demethylation efficiency can be influenced by the space between targeted and methylated sites. To our knowledge, this is the first reported method for targeted demethylation of specific mRNA in transcriptome.

Strikingly, the demethylation seems to be not influenced by located on either 5’ or 3’ side to the dCas13b targeted sites, but it may be dependent on the space between the dCas13b targeted and m^6^A methylated site. Our data showed that the space between 100∼300 nt might enhance the demethylation efficiency (Figure 2 E), which might be the region that partially blocked by nucleosomes and the bound demethylase most efficiently reaches the next available linker RNA region. Consistently, peaks of methylation ∼200 nt in both upstream and downstream of the PAM site has also been observed in DNA methylation using dCas9– Dnmt3a–Dnmt3L methyltransferase ^34^, zinc fingers ^35^ and TALE ^36^ effectors. Our results revealed that methylation up to 300 nt away from the target site can also be demethylated, which was similarly reported in DNA demethylation system using dCas9–peptide repeat and scFv– TET1 catalytic domain ^25^ or dCas9-MQ1 ^37^ fusions. Further, non-additive effect of dm^6^ACRISPR system for RNA demethylation was observed. This may be caused 1) by the less co-transfection efficiency of multiple gRNAs, 2) by a competition of gRNAs for dCas13b-ALKBH5 proteins, or 3) by a steric hindrance when multiple dCas13b-ALKBH5 proteins are bounded on one single transcript, which finally prevents an increase of demethylation efficiency. Our finding was highly reminiscent of previous reports using dCas9-fused p300 to activate gene expression^17^ or dCas9/sgRNA2.0-directed demethylation system for DNA methylation ^38^. Future work will show if improved systems can be developed to overcome these barriers.

Theoretically, the most efficient RNA demethylation should be deposited directly next to the dCas13b binding sites; nevertheless, we observed efficient RNA demethylation for CYB5A by targeting >100 nt away from the methylation sites. We detected limited demethylation effects at two of the predicted, highest scoring off-target regions, which was much weaker than the demethylation at the corresponding on-target sites by same gRNAs. The weak off-target demethylation at near-cognate binding sites might be due to the transient dCas13b binding at these sites, which allows a time window for RNA demethylation deposition. Similar off-target results have been observed in DNA methylation using dCas9–Dnmt3a–Dnmt3L methyltransferase ^34^. It might be reduced by reducing the level of the dCas13b fusion proteins and developing highly specific Cas13b mutants. In addition, untargeted dm^6^ACRISPR or gRNAs did not cause demethylation of the targets nor of their mRNA expressions.

On one hand, our epigenetic editing tools targeting RNA demethylation caused upregulation of mRNA stability of tested gene CYB5A, indicating that dm^6^ACRISPR could be used as a universal tool for gene repression. On the other hand, RNA modification-based therapies could achieve similar changes in genetic information with the advantage of being transient, which eliminates the concern of introducing permanent alterations ^39 40^. Another potential advantage of targeted demethylation is that it is possible to trigger durable effects on specific targets. Notably, the ubiquitous nuclear activity of dCas9-methyltransferases points to an important difference between genetic and epigenetic editing tools, which require unique experimental considerations ^41^. Our future work will modify and optimize our dm^6^ACRISPR system and further explore potential applications, especially in establishing the functional significance of RNA demethylation in the control of the cellular differentiation state. Collectively, considering that CRISPR-based manipulation of the epigenome ^25^ has achieved great success in *in vivo* applications, our developed dm^6^ACRISPR targeting RNA demethylation shows unique potential to correct epimutations in disease states.

## Materials and methods

### Cloning

The original PspCas13b plasmid (Addgene plasmid #103866), gRNA plasmid (Addgene plasmid #103854) and non-targeting gRNA plasmid (Addgene plasmid #103868) were purchased from Addgene. PspCas13b-Alkbh5, Alkbh5-PspCas13b and gRNA-containing plasmids were constructed by Synbio Technologies Company (Suzhou, China). Plasmid containing double mutations at A133H and A1058H of PspCas13b without fusion protein was constructed as active Cas13b.

### Design of the guide RNAs for the CYB5A

Considering that there are multiple mRNA isoforms of CYB5A genes, mRNA sequences of all isoforms were subjected to alignment analysis, and the common regions were acted as targeting candidates for gRNA design. gRNAs targeting 5’UTR, CDS and 3’UTR regions of CYB5A were designed and listed at Table S1, respectively, all designed gRNAs were subject to MEGABLAST (https://blast.ncbi.nlm.nih.gov/Blast.cgi) to avoid mismatching to unexpected mRNA in human genome.

### Mammalian cell culture and plasmid transfection

HEK293T (ATCC) and HeLa (ATCC) cells were cultured in Dulbecco’s Modified Eagle Medium (DMEM, Gibco/Life Technologies) supplemented with 10% fetal bovine serum (FBS, Gibco/Life Technologies), and 1% penicillin/streptomycin (P/S, Invitrogen) under 5% CO_2_ condition. Plasmid transfections were achieved by lipofectamine 3000 (Invitrogen) following the manufacture’s protocol. For 6-well assays, cells were transfected with 1.5 μg PspCas13b-Alkbh5/Alkbh5-PspCas13b/Cas13b and 1μg gRNA for 24 hr before analysis.

### Total RNA isolation and quantitative PCR

After 24hr transfected by Cas13b/PspCas13b-fusion and gRNA plasmids, total RNA was harvested and extracted by Trizol (Invitrogen) according to the protocol used in our previous study ^42^. Extracted total RNA were quantified and reverse transcribed into cDNA using the PrimeScript RT Reagent Kit (TaKaRa). All qPCR reactions were performed as 10 μl reactions using TB Green™ Premix Ex Taq™ II (TaKaRa). Primers for qPCR were listed in Table S2. GAPDH were used as the endogenous control and target C_t_ values were normalized to non-targeting (NT) gRNA C_t_ value. Relative abundance was determined using 2^-ΔCt^. All assays were performed with three independent biological replicates.

### SELECT qPCR

SELECT qPCR method was following Xiao’s protocol ^32^. Briefly, total RNAs were quantified by Qubit (Thermo Fisher Scientific) with Qubit™ RNA HS Assay Kit (Thermo Fisher Scientific). Total RNA (1500 ng) was mixed with 40 nM up primer, 40 nM down primer and 5 μM dNTP in 17 μl 1 x CutSmart buffer (NEB). The RNA and primers were incubated at a temperature gradient: 90°C for 1min, 80°C for 1min, 70°C for 1min, 60°C for 1min, 50°C for 1 min and 40°C for 6 min. Then RNA and primers mixture were mixed with 3 μl of 0.01 U Bst 2.0 DNA polymerase, 0.5 U SplintR ligase and 10 nM ATP, and then incubated at 40°C for 20min and denatured at 80°C for 20min. Afterwards, 20 μl qPCR reaction was preformed, containing 2 μl of final reaction mixture, 200 nM SELECT primers and 2 x SYBR Green Master Mix (TaKaRa). SELECT qPCR program was perform as following condition: 95°C, 5min; (95°C, 10s; 60°C, 35s)×40 cycles; 95°C, 15s; 60°C, 1min; 95°C, 15s; 4°C, hold. Primers for SELECT qPCR were listed in Table S3. C_t_ values of samples were normalized to their corresponding C_t_ values of control. All assays were performed with three independent experiments.

### Immunofluorescent assay

HEK293T cells were seeded onto coverslips in 4-well plate at a density of 1×10^4^ cells per well. After 24 hr transfection of Alkbh5/PspCas13b fusions as indicated, cells were rinsed with PBS and fixed in 4% paraformaldehyde. The cells were permeabilized with 0.1% Triton X-100 in PBS and blocked with 0.1% Tween 20 and 1% horse serum in PBS. Cells were incubated with anti-Alkbh5 antibody (ABE547, Millipore), followed by FITC-labeled Goat Anti-Rabbit IgG (H+L) (A0562, Beyotime). The cell nuclei were stained with Hochest. Localization of Alkbh5 was imaged using Zeiss LSM710 fluorescence microscope.

### LC-MS/MS assay

The mRNA was purified from total RNA using oligo dT magnetic beads. 200 ng purified mRNAs were incubated with nuclease P1 (0.5 U, Sigma) in 25 μl reaction system containing 10 mM NH_4_OAc (pH = 5.3) at 42 °C for 1 h, followed by additions of NH_4_HCO_3_ (1 M, 3μL) and alkaline phosphatase (1 μl, 1 U/μl; Sigma) and incubation at 37 °C for 2 h. After neutralized by 1 μl HCl (3 M), samples were diluted to 50 μl and filtered by 0.22 μm filter (Millipore). All samples (10 μl for each injection) were separated by a C18 column (Agilent) using reverse-phase ultra-performance liquid chromatography and analyzed by an Agilent 6410 QQQ triple-quadrupole LC mass spectrometer using positive electrospray ionization mode. All nucleosides were quantified by use of retention time and ion mass transitions of 268.0 to 136.0 (A) and 282.1 to 150.0 (m^6^A). The quantification was calculated on the basis the standard curves from standards running at the same batch. The ratio of m^6^A to A was calculated based on the calibration curves.

### m^6^A-RIP qPCR

Protein G Magnetic beads were incubated with 1μg m^6^A or IgG antibody in 1x Reaction buffer (150mM NaCl, 10mM Tris-HCl, pH 7.5, 0.1% NP-40 in nuclease free H_2_O) at 4°C for 3hr. 200 μg extracted RNA were added into m^6^A-or IgG-conjugated Protein G Magnetic Beads at 4°C for 3hr. The bound RNAs were incubated with 100 µl Elution Buffer (75nM NaCl, 50 nM Tris-HCl, pH 7.5, 6.25 nM EDTA, 1% (w/v) SDS, 20 mg/ml Proteinase K) for 30min at room temperature. Eluted RNA was recovered with phenol: chloroform extraction followed by ethanol precipitation. m^6^A-RIP RNA was reverse transcribed into cDNA and subjected to qPCR for quantification. IP enrichment ratio of a transcript was calculated as the ratio of its amount in IP to that in the input yielded from same amounts of cells.

### mRNA stability

The HEK293T cells were pre-transfected gRNA and dCas-13b or dCas13b-ALKBH5 constructs for 24 h and then further treated with actinomycin D (Act-D, Catalog #A9415, Sigma, U.S.A) at 5 μg/ml for the indicated timer periods. Cells were collected and RNA was isolated for Real time PCR. Half-life (t1/2) of CYB5A mRNA was calculated using ln2/ slope and GAPDH was used for normalization.

### Statistical analyses

Data was reported as mean ± SD from at least three independent experiments unless otherwise specified. Data was analyzed by two-tailed unpaired Student’s t-test between two groups and by One-Way ANOVA followed by Bonferroni test for multiple comparison. Statistical analysis was carried out using SPSS 16.0 for Windows. All statistical tests were two-sided. \**p*<0.05, \*\**p*< 0.01; NS, no significant.

## Acknowledgements

This research was supported by the National Natural Science Foundation of China (Grant No. 81673454, No. 81672608, No. 31801197, and No. 81572270), the Guangdong Natural Science Funds for Distinguished Young Scholar (No. 2014A030306025), the Guangdong Province Key Laboratory of Malignant Tumor Epigenetics and Gene Regulation (2017B030314026), and the Fundamental Research Funds for the Central Universities (Sun Yat-sen University) (No. 16ykpy09). We thank Prof Dali Li at East China Normal University and Dr Guifang Jia at Peking University for discussion and technical support.

