## Supplementary material for "Targeted mRNA demethylation using an engineered dCas13b-ALKBH5 fusion protein": Figure S1~S2, and Table S1 to S3

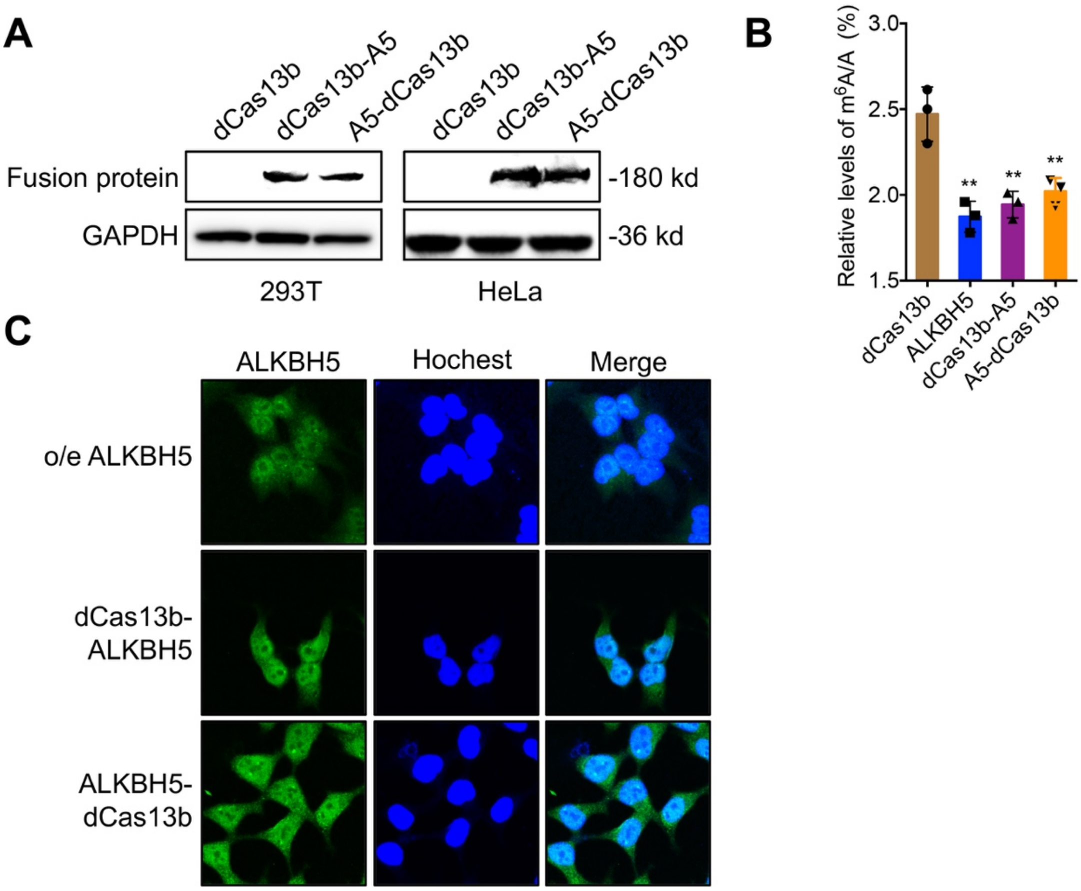


**Figure S1 In vivo function and localization of dCas13b-ALKBH5.**

(A) The fusion protein dCas13b-ALKBH5 or ALKBH5-dCas13b in transfected cells was measured by western blot with anti-ALKBH5 as primary antibody;

(B) HEK293T cells were transfected with vector control (dCas13b), ALKBH5, dCas13b-ALKBH5, or ALKBH5-dCas13b for 24 h, the $m^6A/A$ ratio of total mRNA were determined by LC-MS/MS;

(C) After transfected with ALKBH5 construct, dCas13b-ALKBH5, or ALKBH5-dCas13b for 24 h, the subcellular localization of fusion protein was checked by confocal using an antibody against ALKBH5.

Data are presented as the mean ± SD from three independent experiments. ** $p<0.01$ compared with control.

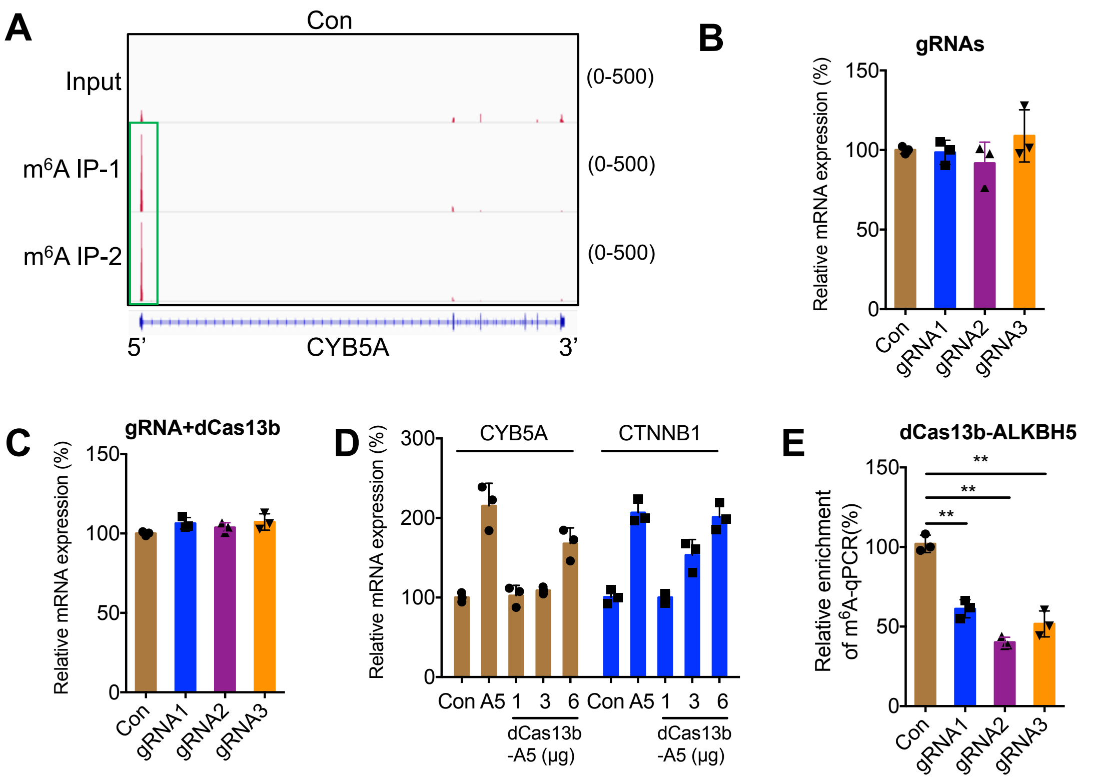


**Figure S2 The dm$^6$ACRISPR induced m$^6$A demethylation at CDS increases mRNA stability.**

(A) m$^6$A peaks were enriched in CDS of CYB5A genes from m$^6$A RIP-seq data. Squares marked increases of m$^6$A peaks in cancer cells;

(B) The expression of CYB5A in HEK293T cells transfected with gRNAs for 24 h;

(C) The expression of CYB5A in HEK293T cells transfected with dCas13b and gRNAs for 24 h;

(D) Cells (1X10$^6$) were transfected with vector control (dCas13b), ALKBH5 construct (1μg), or increasing amount of dCas13b-ALKBH5 for 24 h, the mRNA expression was checked by qRT-PCR;

(E) m$^6$A RIP-qPCR analysis of CYB5A mRNA in cells transfected with dCas13b-ALKBH5 combined with gRNA control or gRNA1/2/3, respectively, for 24 h.

Data are presented as the mean ± SD from three independent experiments. ** $p<0.01$ compared with control.

Table S1 The sequences of gRNA used in this study

| Name | Guide RNA sequence |
|---|---|
| gRNAcyb5a-1 | AGCAGAGCGCGCGACTCAGCCAGCTCCACCCGGGACATTCGT TGTGGAAGGTCCAGTTTTGAGGGGCTATTACAAC |
| gRNAcyb5a-2 | CTGGATGGAGCTCCCCAATGATGAATGTTTTGGACATTTCGTT GTGGAAGGTCCAGTTTTGAGGGGCTATTACAAC |
| gRNAcyb5a-3 | TGTCCAAAGCAGGCTCTTCCTGCGCTGACTTCTGAGGAGGGT TGTGGAAGGTCCAGTTTTGAGGGGCTATTACAAC |

Table S2 Primers for PCR

| Gene | Forward | Reverse |
|---|---|---|
| Plasmid mutation | | |
| Cas13b-A133H | CAGGGACCTGACCAACCACTAC AAGACCTACGA | TCGTAGGTCTTGTAGTGGTTGGT CAGGTCCCTG |
| Cas13b-A1058H | CCGGAACGCCTTCGATCACAAC AATTACCCCGA | TCGGGGTAATTGTTGTGATCGAA GGCGTTCCGG |
| qRT-PCR | | |
| GAPDH | GTCTCCTCTGACTTCAACAGCG | ACCACCCTGTTGCTGTAGCCAA |
| CYB5A | GTTTTAAGGGAACAAGCTGGAG | TCCACCAACTGGAACTAGAATC |
| GRSS56 | GCTGTGCTGCTGCTGCTACC | CTGCCTGCAACGCCTGAGTC |
| GALR1 | TTCCTTCCTCTTCAGAATCACC | CGAATGTGACACTTGAACACTT |
| INPP4A | GAATCCGATCCAAATACGCTTC | CTCTAGTAGGAGCTTCACGAAC |

Table S2 Primers for SELECT qPCR

| Name | Sequence |
|---|---|
| CYB5A-X-up | TAGCCAGTACCGTAGTGCGTGGTGGTTGTGCTTCTGAATC |
| CYB5A-X-down | 5PHOS/CCTCTAGGGTGTAGTACTTCCAGAGGCTGAGTCGCTGCAT |
| CYB5A-N-up | TAGCCAGTACCGTAGTGCGTGTTCTGAATCTCCTCTAGGG |
| CYB5A-N-down | 5PHOS/GTAGTACTTCACGGCCTCGTCAGAGGCTGAGTCGCTGCAT |
| qPCR-SELECT-Forward | ATGCAGCGACTCAGCCTCTG |
| qPCR-SELECT-Reverse | TAGCCAGTACCGTAGTGCGTG |
